# Tissue-engineered collagenous fibrous cap models to systematically elucidate atherosclerotic plaque rupture

**DOI:** 10.1101/2021.07.20.451997

**Authors:** T.B. Wissing, K. Van der Heiden, S.M. Serra, A.I.P.M. Smits, C.V.C. Bouten, F.J.H. Gijsen

## Abstract

A significant amount of vascular thrombotic events is associated with rupture of the fibrous cap that overlie atherosclerotic plaques. Cap rupture is however difficult to predict due to the heterogenous composition of the plaque, unknown material properties, and the stochastic nature of the event. Here, we aim to create tissue engineered human fibrous cap models with a variable but controllable collagen composition, suitable for mechanical testing, to scrutinize the reciprocal relationships between composition and mechanical properties.

Myofibroblasts were cultured in 1 × 1.5 cm-sized fibrin-based constrained gels for 21 days according to established (dynamic) culture protocols (i.e. static, intermittent or continuous loading) to vary collagen composition (e.g. amount, type and organization). At day 7, a soft 2 mm Ø fibrin inclusion was introduced in the centre of each tissue to mimic the soft lipid core, simulating the heterogeneity of a plaque.

Results demonstrate reproducible collagenous tissues, that mimic the bulk mechanical properties of human caps and vary in collagen composition due to the presence of an successfully integrated soft inclusion and the culture protocol applied. The models can be deployed to assess tissue mechanics, evolution and failure of fibrous caps or complex heterogeneous tissues in general.

## Introduction

Atherosclerosis is characterized by the accumulation of lipids within the intimal layer of the arterial wall, forming so-called plaques (Fig.1A), that upon rupture can cause acute and often lethal manifestations like stroke or myocardial infarction^1,2^. Although mortality rates significantly decreased in the last decades, an atherothrombotic event remains one of the leading causes of mortality globally^3,4^. Various stages of plaque development can be identified. Thin cap fibroatheromas (TCFAs) are most prone to rupture and are therefore termed “vulnerable” plaques.^2^ Vulnerable plaques are characterized by a soft lipid core embedded in a collagenous matrix, separated from the bloodstream by a thin, often inflamed and calcified, collagenous cap (i.e. the fibrous cap) (Fig.1B).^5^ Current clinical initiatives to detect vulnerable plaques predominantly focus on non-invasive imaging modalities to visualize plaque features that are associated with the mechanical stability of the plaque and could therefore be indicative of rupture risk^6^, like cap thickness^7^, the size of the lipid core^8^ and the presence of inflammatory cells^9^, calcifications^10^ and intraplaque hemorrhages^11^. However, recent prospective clinical studies revealed a poor predictive value of these vulnerability features for clinical events ^12^.

**Fig.1.**
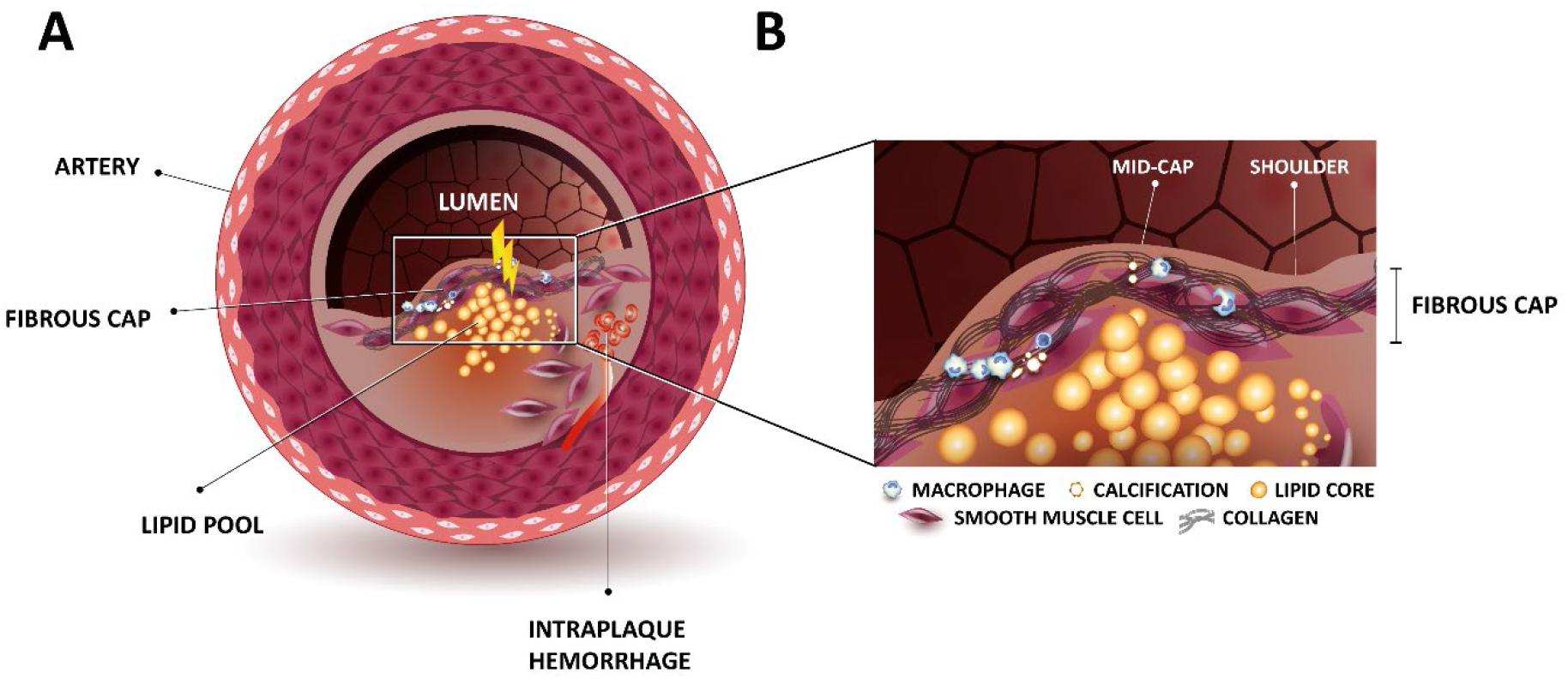
Simplified graphical representation of a cross sectional view of a vulnerable plaque that constitutes of an intraplaque hemorrhage and a lipid core **(A)** sealed by an collagenous fibrous cap **(B)**.

The stability of a plaque largely depends on the integrity of the fibrous cap, but little is known about its mechanical properties and rupture mechanism. It is however suggested that the locus of fibrous cap rupture might be related with physical (in)activity. More specifically, it was observed that plaques that rupture during physical rest predominantly rupture at the shoulder region, while plaques that rupture during physical activity predominantly fail at mid cap section (Fig.1B to clarify these cap loci).^13^ Although these observations suggest different rupture mechanisms, the underlying principle remains the same: the cap ruptures when mechanical loads exceed local tissue strength.

Essential for tissue strength is the fibrillar collagenous matrix, which is the main load-bearing structure of the fibrous cap. The matrix owes its mechanical properties to a combination of factors, like collagen content (i.e. amount, type and crosslinking), collagen fiber distribution and the presence of degrading compounds (e.g. matrix metalloproteinases (MMPs)) that locally damage the collagenous matrix^14–17^. The mechanical properties of the collagenous matrix are subject to change, as production, remodeling, and degradation are known to be locally controlled by the mutual interplay between the cellular population (e.g. vascular smooth muscle cells (VSMCs) and immune cells) and environmental cues, like mechanical strain ^18–20^.

The fibrous cap predominantly consists of fibrillar collagen type I and III ^21–23^. How collagen fibers are distributed varies throughout and between caps, though Johnston et al. suggested that circumferentially orientated collagen fibers in the cap play the most dominant role for determining plaque stability.^15,24^ The strength and stability of these fibers is tightly regulated by intra- and intermolecular cross-links that are generated during collagen maturation. Lysyl oxidase (LOX) mediated enzymatic crosslinking is one of the main extracellular processing steps that impacts collagen tensile strength and its resistance to proteolytic enzymes. LOX is a copper-dependent enzyme, secreted by plaque resident cells, including VSMCs and myofibroblasts, that catalyzes the formation of the covalent intermolecular mature cross-links hydroxylysylpyridinoline (HP) and lysylpyridinoline (LP). HP and LP cross-link formation results in the stabilization of extracellular collagen, making it less susceptible to proteolytic degradation.^25–28^ Simultaneously, extracellular glycation- or oxidation-induced crosslinking, also called advanced glycation end products (AGEs)-related crosslinking, impairs functional interaction of collagen with the cellular components and stiffens the matrix, making it more brittle^26,29–31^. It predominantly occurs within ageing and diabetic patients, two major patient groups at risk of acute manifestations of atherosclerosis. Additional mechanisms behind reduced collagen strength include the local production of MMPs and cathepsins, predominantly by infiltrated immune cells like macrophages, that cleave and induce local weakening of the collagenous matrix, reducing cap strength^32–35^.

Various cap strength values are reported in literature, ranging from 158 kPa^36^ to 870 kPa^24^. The variability in reported strength values is most probably related to variances in cap composition and loading, affecting the locus of peak mechanical stress and eventual failure mechanism^24^. Correspondingly, several plaque failure mechanisms have been proposed, including delamination^37^, debonding^38^, cavitation^39^, fatigue^40^ and failure^35^. However, due the paucity of experimental data, the conditions that underlie each failure mechanism remain still largely unknown, complicating fibrous cap failure predictability.

Understanding rupture critically depends on systematically analyzing the relationships between i) the biological plaque and cap components, ii) the mechanical strength of these components, and iii) the rupture mechanism^41^. Unfortunately, adequate monitoring of such interacting plaque features in patients is unethical and the stochastic nature of the event limits researchers to analyzing postmortem plaque material. The *ex vivo* mechanical investigation of already ruptured plaques is complex due to the destroyed morphology. Moreover, plaques that did not rupture are very heterogeneous in composition and potentially more stable and as such not fully representative for the rupturing plaque. As an alternative, animals models are frequently used but these are limited as plaque development and cap rupture remain stochastic and these animal models do not adequately mimic human plaque development and mechanics^41^. Innovative models should be developed that aid in the systematic investigation of human plaque rupture with high experimental control in a relatively high throughput fashion. Within this study, we take the first step in simplifying and controlling plaque complexity, starting with the fibrous cap and its load-bearing collagenous matrix.

It was shown in the field of cardiovascular tissue engineering that collagen matrix composition (i.e. amount, type, crosslinking and organization) can be manipulated by tuning the cell culture conditions ^17,19,42–46^. Based on these insights, we evaluate three (dynamic) culture strategies to obtain tissue engineered human fibrous cap models that vary in collagen organization and content in a controlled manner, and approach the bulk mechanical properties reported for human caps. More specifically, myofibroblasts seeded fibrin-based constrained gels were exposed to either 1) an intermittent straining protocol (I-strain), with alternating periods of straining and rest, to promote collagen synthesis and remodeling (i.e. evoking fiber anisotropy), or 2) a continuous cyclic straining protocol (C-strain) to shift this balance to collagen remodeling and maturation, stimulating alignment and crosslinking, improving tissue mechanical properties ^17,19,46–48^. As a third group, statically cultured samples were included to obtain more isotopically orientated collagenous tissues. Of note, a soft inclusion (i.e. lipid core analog) was integrated in each model type, as it is known that local variations in matrix stiffness create heterogeneous stress and strain distributions that dictate collagen formation and remodeling ^45^.

## Materials and Methods

### Experimental outline

Figure 2 provides a graphical representation of the *in vitro* fibrous cap model (Fig.2A), the (dynamic) cell culture protocols and the analyses performed (Fig.2B). In brief, vascular derived myofibroblasts (i.e. human vena saphena cells (HVSCs)) were seeded in fibrin-based gels, longitudinally constrained between two Velcro strips (1 x 1.5 cm-sized fibrin gels). After 7 days of static culture a 2 mm Ø core was punched in the center of each sample and filled with fibrin to form a soft inclusion as an analog for the lipid core. It was hypothesized that local variations in matrix stiffness, due to the integration of a soft inclusion, albeit in combination with dynamic loading would result in variations in the local collagen architecture ^45^. The samples were statically cultured for another week to ensure integration of the soft inclusion within the tissue. Next, the samples were equally divided amongst three experimental groups and exposed to either dynamic loading (intermittent −1hr 4% strain followed by 3hrs of rest- or continuous −4%-straining) or no loading (static) up to day 21. At day 21, samples were exposed to uniaxial tensile tests and further analyzed with tissue assays, histological analyses and imaging to assess mechanical properties and collagen composition (i.e. amount, type, orientation and crosslinking), respectively.

**Fig.2.**
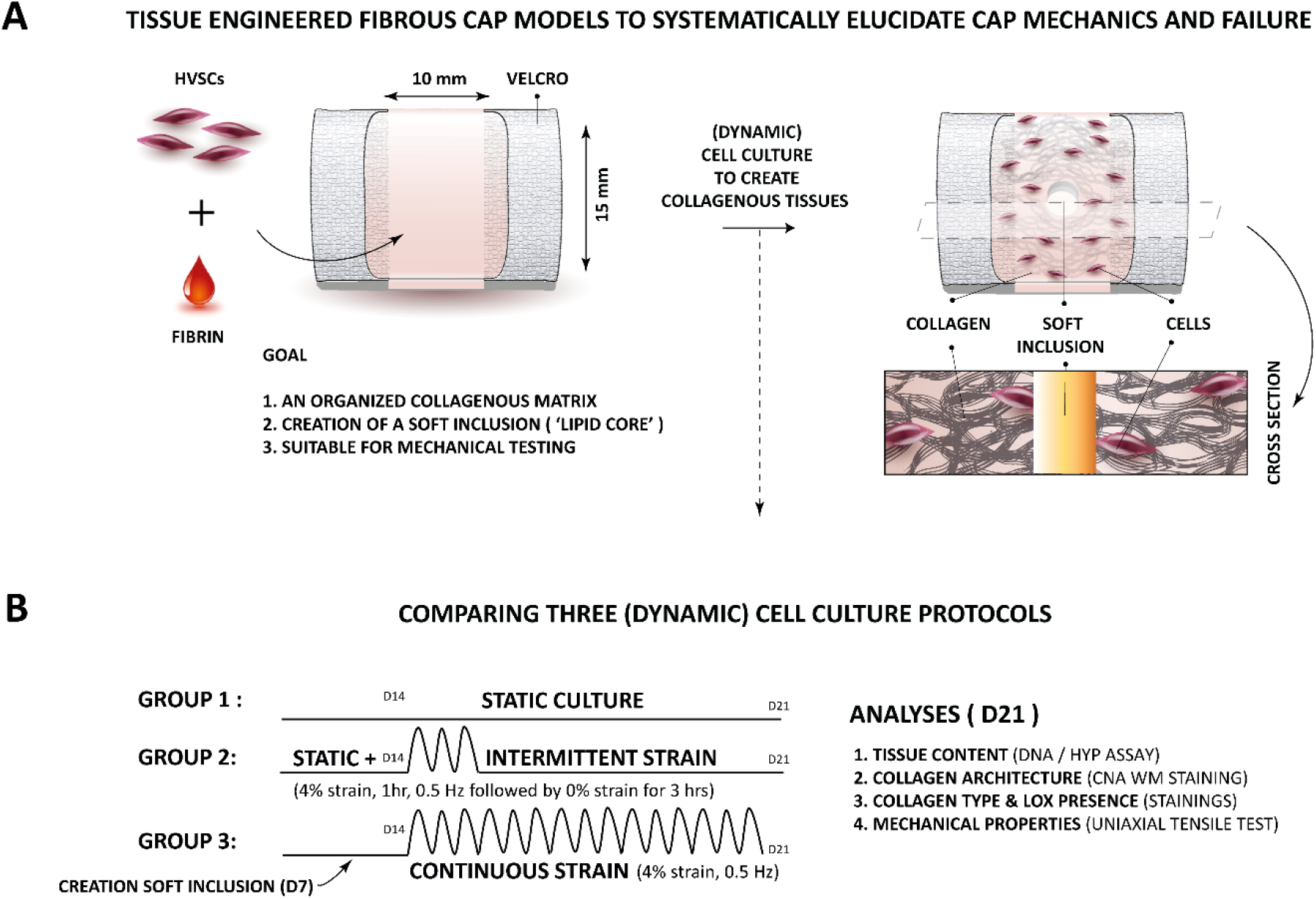
Graphical representation of the tissue engineered *in vitro* fibrous cap models generated. Collagenous tissues were generated by seeding HVSCs in fibrin gels between pares of Velcro strips and exposing them to three different (dynamic) cell culture protocols **(A)**. More specifically, samples were first statically cultured for 14 days, whereafter they were exposed to another 7 days of static culture (group 1), 7 days of intermittent strain (group 2, i.e. alternating periods of straining (1hr, 4% strain) and rest (3hrs)) or continuous strain (group 3, i.e. 4% strain). A soft inclusion was created at day 7 to mimic plaque heterogeneity. At day 21, samples were analyzed to determine cell number and collagen content, collagen architecture, collagen type, LOX presence and mechanical properties **(B)**. Abbreviations: human vena saphena cells (HVSCs), hydroxyproline (HYP), whole mount (WM), lysyl oxidase (LOX).

### Engineering of fibrin-based tissues

#### Mold preparation

Rectangular Velcro strips of 5×15 mm each were attached to the flexible membranes of 6-well Bioflex culture plates (untreated, Flexcell Int, McKeesport, PA) (dynamic loading groups) or the bottom of normal tissue culture well plates (static control) using medical adhesive silicone (Silastic MDX4-4210, Dow Corning, Midland, MI). Per well, two Velcro strips were placed in parallel 10 mm apart to form the base of the mold. These Velcro strips allow cells to grab onto, while simultaneously providing a grip for mechanical testing. Silicon glue to attach the Velcro strips was dried overnight in an oven at 60°C whereafter the strips were incubated for 30 minutes with 5 ml of 70% EtOH/well at room temperature (RT). After this incubation period, the wells were rinsed with sterile PBS (3x 5 min) and placed under UV for 30 minutes to decontaminate.

Rectangular polydimethylsiloxane (PDMS) bars (10×4×4 mm) were created by mixing a silicon elastomer base and a curing agent (10:1 ratio) and curing the mixture overnight at RT. The bars were sterilized using 70% EtOH, washed with sterile PBS (3x 5 min) and placed on the sides in between the Velcro strips to create a rectangular mold (dimensions: 10×15 mm) for the fibrin-cell suspension to solidify during the initial phase of the culturing process. The bars were removed on day 5 of cell culture.

#### Human vena saphena cell culture

HVSCs were isolated from the human vena saphena magna of a 63-year old women after coronary bypass surgery according to previously established protocols^49^ conform the Dutch advice for secondary-use material. The cells were cultured in a standard incubator (37°C, 5% CO_2_) in complete growth medium consisting of advanced-DMEM supplemented with 10% Fetal Bovine Serum, 1% Glutamax and 1% Penicillin/Streptomycin. Cells were passaged at 80% confluency and used for the experiment at passage 7.

### Preparation of collagenous tissues

To create the tissues, HVSCs were seeded in a fibrin matrix that served as a temporary matrix for cells to deposit their own extracellular matrix (ECM). A suspension of bovine fibrinogen (10 mg/ml, Sigma F8630), bovine thrombin (10 U/ml, Sigma T4648) and HVSCs (1.5×10^6 cells/ml) were mixed and seeded in the mold between the Velcro strips and the PDMS bars, as well as covering half of each Velcro strip to allow cells to anchor to the Velcro brushes. Potential air bubbles were removed before solidification of the fibrin gel, using a sterile needle (BD Microlance, 24G). The samples were placed in an incubator (37°C, 5% CO_2_) for 30 minutes to gelate, whereafter L-ascorbic acid 2-phosphate (vitamin C, 0.25 mg/ml) and ε-Amino Caproic Acid (ε-ACA, 1 mg/ml) enriched medium (5 ml/well) was added. Vitamin C and ε-ACA were added to the medium to stimulate collagen synthesis and reduce fibrin degradation, respectively. Medium was refreshed every 3 days with ε-ACA being added during the first 7 days, to prevent fibrin break-down and allow cells to deposit their own ECM ^45^.

### Creating a soft inclusion to mimic the soft content of the lipid core and heterogeneity of a plaque

After 7 days of static cell culture, media were removed and a core (2 mm Ø) was punched in the middle of each tissue with a sterile biopsy punch (Kai Medical, Japan). The circular tissue was removed and filled with a suspension of bovine fibrinogen (10 mg/ml, Sigma F8630) and bovine thrombin (10 U/ml, Sigma T4648) to form a fibrinogenic soft inclusion (stiffness 0.5-1.0 kPa)^50^ that represents the lipid core region. The tissues were placed in a standard incubator (37°C, 5% CO_2_) for 15 minutes to allow the suspension to solidify, whereafter fresh vitamin C enriched medium was added.

### Dynamic cell culture to stimulate collagen formation and remodeling

The fibrin-based gels in the Bioflex plates were placed on the Flexcell FX-40001 (Flexcell Int, McKeesport, PA). The Flexcell system applies a vacuum onto the compartment under the flexible silicon membrane pulling the membrane over a post to induce a strain in the gel. The samples were exposed to either 7 days of I-strain (4% strain for 1 hr (0.5 Hz) followed by 3 hrs of rest) or C-strain (4% strain, 0.5 Hz) to promote collagen synthesis and remodeling (i.e. evoking fiber anisotropy), or collagen remodeling and maturation, respectively ^17,19,46–48^.

#### Strain Validation

In order to validate the amount of strain applied to the tissues during culture (set to 4% in the software), Digital Image Correlation (DIC) analysis was performed on the images of maximum compaction and extension in the direction of loading. Videos were captured at day 21 and converted to images at 30 Hz in MATLAB (Mathworks, Massachusetts, USA), whereafter the images of maximum compaction and extension were selected. Subsequently, the average strains in the y direction (ε_yy_) per loading regime group were calculated (supplementary Tab.S1) in the region of interest (ROI) (i.e. mid-cap section, supplementary Fig.S1) using the open source 2D DIC software Ncorr^51^ (see paragraph DIC).

#### Tissue analyses

At day 21, tissues were processed for further analysis. Samples were either 1) washed in PBS (3x 5 min), fixated for 1 hr in 3.7% formaldehyde, followed by a PBS washing step (3x 5 min) and stored at 4° Celsius, or 2) directly used for uniaxial tensile testing. After the uniaxial tensile test, left-over material was washed in PBS, snap frozen in liquid nitrogen and stored at −80°C. At a later timepoint, these samples were used to determine DNA, hydroxyproline (HYP) and crosslinking content. Fixated samples were used for histological analysis to determine collagen orientation, collagen type and the presence of LOX.

### Tissue composition

The tissues that were reserved for the DNA, HYP and crosslinking determination were consecutively lyophilized, weighted and digested in papain enriched digestion buffer (100 mM phosphate buffer (pH=6.5), 5mM L-cysteine (C-1276), 5mM ethylene-di-amine-tetra-acetic acid (EDTA, ED2SS), and 140 μg/mL papain (P4762), all from Sigma-Aldrich). Each sample was cut in small pieces, mixed with 300 μl of digestion buffer in a new Eppendorf tube and placed at 60°C (16 hrs, Thermormixer Compact F 1.6 A,VWR) for digestion. After digestion, samples were stored at 4°C until analyses. Prior to DNA quantification, samples were vortexed and centrifuged (12,000 rpm, 10 minutes on 4°C) to mix and precipitate remnants, respectively. DNA content was quantified using 10 μL of the supernatant, the Qubit dsDNA BR assay kit (Life Technologies, Carlsbad, California, USA) and the Qubit fluorometer (Life Technologies) following manufacturer’s instructions and normalized to tissue dry mass. To quantify HYP content digested samples were first hydrolyzed using 16 M sodium hydroxide (Merck; B1438798). Subsequently, HYP content was quantified with a Chloramin-T assay, including trans-4-hydroxyproline as a reference (Sigma; H5534).^52^ HYP values were normalized to total dry mass and DNA content to assess overall collagen formation and collagen production per cell.

### Bright field microscopy

Brightfield microscopy images were taken of each soft inclusion after one and two weeks of culture to assess cellular infiltration and soft inclusion geometry and integrity (i.e. no visible defects within the soft inclusion or at the tissue-soft inclusion interface). Soft inclusion geometry was quantified using the Image J software (U.S. National Institutes of Health, Bethesda, MD, USA)

### Matrix content and organization

Whole-mount samples or 7 μm thick cryosections were analyzed to evaluate matrix content and organization using either fluorescence stainings, histology or immunohistochemistry (IHC). As a positive control, 7 μm thick cryosections of a human atherosclerotic plaque of a 68 year old male were included. This carotid endarterectomy specimen was retrieved in line with the ethical guidelines of the 1975 Declaration of Helsinki, after written informed consent from the patient and approval of the Medical Ethics Committee of the Erasmus Medical Center. After endarterectomy, the plaque specimen was snap frozen, stored at −80 °C and embedded, similar as the collagenous tissue engineered fibrous cap models, in Tissue Tek, to allow sectioning.

#### Collagen organization

Whole-mounts samples were incubated with an Oregon Green labeled CNA35 probe (CNA35-OG488) and the nuclear stain DAPI to assess collagen and cellular organization. The in house developed CNA35 probe binds to a variety of fibril forming collagens, including type I and III, with little to no specificity for other extracellular matrix proteins ^53^. The probe was diluted in PBS to a concentration of 1 μM and incubated together with DAPI (1:500, Sigma) for 16 hrs at a shaker (300 rpm) at RT. The samples washed in PBS (3x 5 min) and embedded in Mowiol prior to confocal laser scanning microscope analysis (Leica TCS SP8X). Collagen fiber organization was determined as described in the supplementary information (supplementary Fig.S2-4 and supplementary Eq.S1.).

#### Matrix morphology and composition

Fibrillar collagen or cell nuclei and cytoplasm were visualized in cross-sections stained for Picrosirius Red (PSR) or Hematoxylin and Eosin (H&E), respectively. The stained cross-sections were embedded in enthalan and assessed with brightfield microscopy.

#### Collagen type and LOX presence

Formalin-fixed cryosections were immunohistochemically stained to assess the presence of collagen type I (1:200, Sigma), type III (1:250, Abcam) and LOX (1:100, Novus Biologicals). Sections were rinsed in PBS and incubated with 5% (w/v) goat serum in a PBS solution containing 1 % (w/v) Bovine Serum Albumin to block non-specific binding. Primary antibodies (1:100/200/250 in 10× diluted blocking solution) were incubated overnight at 4 °C, followed by a PBS washing step (3×5 min) and another 1 hr incubation step with the fluorescently labeled secondary antibodies (1:500 in 10× diluted block solution, Molecular Probes) at RT. The secondary antibodies were washed away with PBS (3x 5 min) whereafter 4’,6-diamidino-2-phenylindole (DAPI, 1:500, Sigma) was added for 10 minutes to localize the cell nuclei. The samples were mounted in Mowiol and visualized with an inverted fluorescent microscope (Leica DMi8).

### Tissue mechanical characterization

Uniaxial tensile tests were performed at day 21 to assess the mechanical properties and rupture behavior of the samples of the different experimental groups using a test set-up equipped with a 5N load cell (CellScale Biomaterial Testing, Waterloo, Canada). Before testing, samples were removed from the well-plate, washed in PBS and photographed on a grid background to determine the compaction rate in the horizontal (x, sample width) and vertical (y, sample length) direction (Figure 3B). The compaction in the z direction (thickness) was measured using a digital microscope (Keyence VHX-500FE, Itasca, IL).

**Figure 3.**
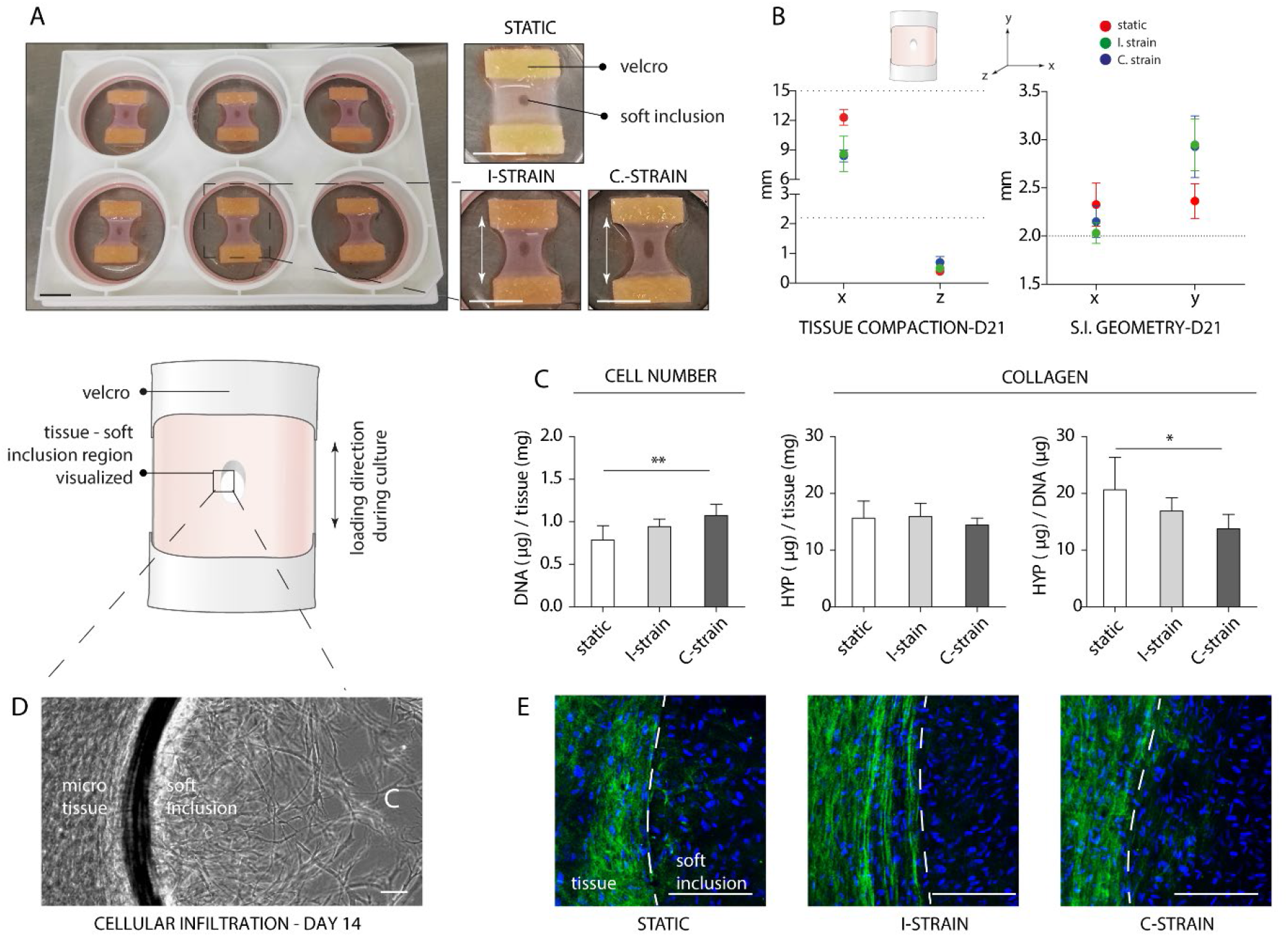
Comparison of three tissue-engineered fibrous cap models at day 21. Representative photographs underlining the reproducibility and rectangular/dog bone shape morphology of the created tissues at day 21. **(A)**. Quantified tissue and soft inclusion geometry at day 21 in comparison to the initial sample and soft inclusion size (dotted lines), respectively (n ≥ 5 per group) **(B)**. Cell number (DNA) and collagen (HYP) levels normalized to total tissue mass and cell content (n ≥ 5 per group) **(C)** Representative bright field image of the tissue-soft inclusion interface before dynamic loading (day 14) demonstrating gradual cellular infiltration. The C indicates the center of the soft inclusion **(D)**. Representative collagen (green) and nuclei (blue) stained zooms of the tissue-soft inclusion interface of each experimental group at day 21 displaying the differences in collagen presence between the artificial fibrous cap (i.e. tissue) and soft inclusion **(E)**. Scale bars, 1 mm (A) and 100 μm (D and E).

The Velcro’s on each side of the sample were placed in the clamps of the tensile tester. Each sample was mounted in the set-up and exposed to uniaxial loading in PBS at 37° Celsius to simulate physiological conditions. After preconditioning (10 cycles of 10% strain^54^), the samples were uniaxially strained at a 100%/min strain rate until failure. A camera and lens were located directly above the tissues, capturing images during testing at a speed of 15Hz for digital image analyses.

The nominal engineering stress-strain curves were derived from the force and displacement measurements utilizing the average initial thickness and width measurements in the ROI (supplementary Fig.S1), assuming incompressibility and plane-stress conditions

Data averaging was performed on the stress-strain data and the stiffness values were computed as the linear slopes at 2%, 5% and 10% strain (i.e. ε_2%_, ε_5%_, ε_10%_,) based on the DIC analyses described below. ε_10%_ was chosen as the upper limit because it represent the maximum amount of physiological circumferential strain experienced by the mid cap region in a fibrous plaque ^55^.

Before mounting the sample in the uniaxial tensile set-up, graphite particles were applied onto the surface of each sample with a cotton swab to facilitate optical strain analysis of the region of interest via digital image correlation. 30% of the sides of each tissue were excluded from analyses to exclude boundary artifacts caused by the clamping and compaction.

DIC analyses were performed as a normalization method to exclude potential clamping and compaction artifacts and allow a consistent comparison between the different samples. The DIC analyses were performed in MATLAB on the ROI using the open source 2D DIC software Ncorr^51^. Local strains were assessed in the ROI to determine at which image there was an average of 12% strain in the direction of loading (ε_yy_). The regions on each side of the soft inclusion were treated separately (supplementary Fig.S1) as local variation may occur. The stress-strain data was scaled according to this image and the averages were calculated (i.e. average between 0-2% strain resulted in an average data point at 1% strain). The linear slope was calculated between 1% and 3% strain, 4% and 6%, and 9% and 11%, in order to extract the stiffness values at 2%, 5%, and 10%, respectively. The average stress-strain curves for each group (static control, I-strain, and C-strain) were extracted, including the standard deviations, to provide a representative plot of the stress-strain behavior for all tissues.

### Statistical analysis

All data (DNA, HYP, compaction and stiffness values) are presented as the mean ± standard deviation unless stated otherwise. Statistical analyses were performed using Prism (GraphPad, La Jolla, CA, USA). A Shapiro-Wilk test and Levene’s test were performed for normality and equal-variances. Statistical significant differences between the experimental groups, in case of proven normality, were detected using an one-way ANOVA with subsequent Tukey’s post-hoc test (assuming equal variances) or Dunnett’s post-hoc test (unequal variances). In case of not normally distributed data, a nonparametric Kruskal-Wallis test with a Dunn’s multiple comparison test was performed. Differences were considered statistical significant for p values < 0.05 (visualized as * p < 0.05; ** p < 0.01; *** p < 0.001).

## Results

### Three reproducible fibrous cap models with integrated soft inclusions

After 21 days of cell culture, consistent tissues were formed between the Velcro strip constraints for each (dynamic) cell culture protocol applied (Fig.3A). Exposure to cyclic strain during the last week of culture resulted in enhanced compaction and consequent dog bone shaped tissues for both the I-strain and C-strain group. Whereas the statically cultured controls exhibited relative low percentages (−18 ± 6%) of compaction in width, higher rates of compaction were seen when loading during cell culture was applied (I-strain: −44 ± 4% and C-strain: −42 ±12%). Smaller differences in compaction in the z direction (i.e. tissue thickness) were observed between the experimental groups, with an overall average thickness of approximately 537 ± 169 μm (Fig.3B).

To quantify collagen content in the tissues, the amount of HYP, a matrix component for stabilization of the collagen molecule, was quantified (Fig.3C). In general, negligible differences in HYP presence per tissue dry mass were found. However, a statistically significant increase in cell number upon exposure to continuous loading in comparison to the static control (p = 0.008) was observed. This increased cell number was accompanied by a significant decrease in amount of HYP presence per cell between the static and continuous strain group (p = 0.013).

At day 7 of culture, soft inclusions were created to mimic the soft lipid core, overlying fibrous cap and heterogeneity of a plaque. With time, the soft inclusions were integrated into the tissues, as characterized by cell and collagen ingrowth into the inclusions (Fig. 3D-E). At day 21, infiltrated cells covered the complete soft inclusion area for all groups assessed, though negligible amounts of collagen were deposited throughout this soft inclusion area in comparison to the rest of the sample(Fig.3E).

### Collagenous tissues with (dynamic) culture protocol dependent collagen organizations

The local collagen fiber dispersion was assessed and quantified (Fig.4).Overall, a more condensed ± 5 - 50 μm thick collagen layer (1 - 10% of the tissue thickness) was detected at the upper surface of each tissue. This densely packed layer demonstrated a clear anisotropic collagen organization, in the direction of the Velcro constraints, irrespective of the type of tissue or locus assessed (supplementary Fig.S5). Throughout the other ≥ 90 percent of the tissue thickness the collagen fiber dispersion was consistent though dependent on the locus analyzed and culture protocol applied. Symmetry in collagen fiber distributions were observed between the top and base and the left and right with respect to the soft inclusion of all tissues within one experimental group. As such, only the results of one quartile per tissue type are presented (Fig. 4).

**Figure 4.**
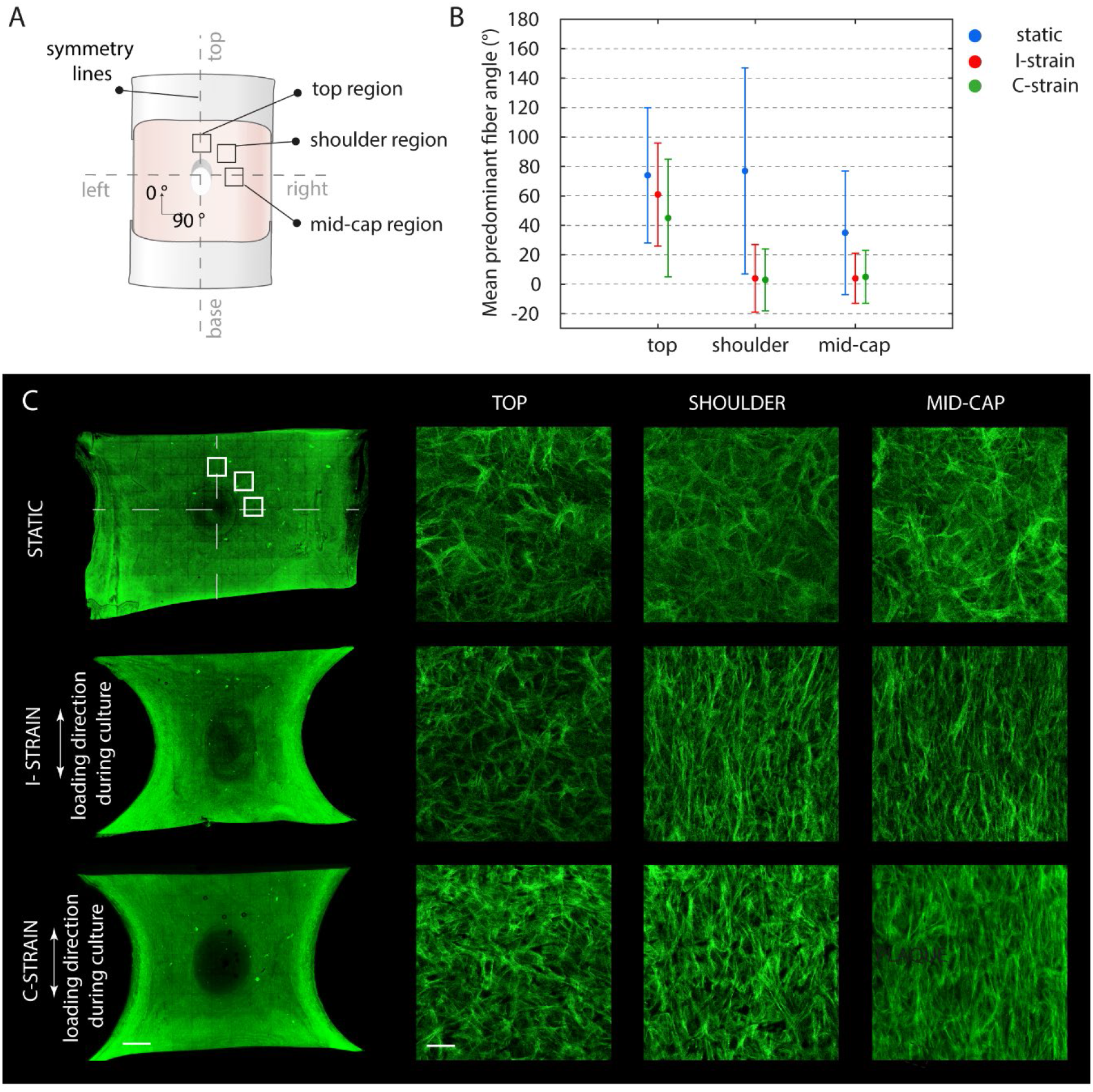
Internal collagen architecture at day 21. Graphical representation of the loci visualized **(A).** Quantification of the collagen fiber distribution (i.e. predominant fiber angles and presence of (an)isotropy). Large SD indicates greater isotropic behavior surrounding mean predominant fiber angle. Small SD indicates greater anisotropic behavior surrounding mean predominant fiber angle **(B)**. Representative confocal tile scans and zooms of CNA35 (a collagen binding protein ^53^, green) stained samples of the same top, shoulder and mid-cap regions as evaluated in B **(C)**. See supplementary Fig.S5 for an overview of the upper superficial layer collagen architecture (first 5 −50 μm of the upper surface of each tissue). Scale bars, 3 mm (tile scans) and 100 μm (zooms).

All top regions analyzed exhibited isotropic collagen organizations irrespective of the culture protocol applied. The predominant fiber angles ranged between 74°degrees for the statically cultured samples up till 61° and 45° degrees when exposed to intermittent and continuous loading, respectively. Moving towards the shoulder and mid cap regions, the fibers appear more anisotropic in the direction of the Velcro constraints for the dynamically loaded samples. Those samples showed predominant fiber angles of 4° and 3° degrees in the shoulder region and 4° and 5° degrees in the mid cap region for the I-strain and C-strain respectively. The static control demonstrated a more isotropic collagen fiber distribution with a predominant fiber angle of 77° degrees in the shoulder and 35° degrees in the mid cap region. Fig. 4 demonstrates the created organized collagenous tissues with increased collagen alignment in the direction of the Velcro constraints when moving from the top to the shoulder and ultimately the mid-cap region, especially when loading was applied during culture.

### LOX positive collagenous tissues that encompass collagen type I and III

Histological assessment of the tissue engineered fibrous cap models tissue content, demonstrated a similar collagenous architecture as observed in the fibrous cap of human atherosclerotic plaques, especially when loading was applied during culture (Fig.5A,B). H&E and PSR staining shows a similar morphology, although the intensity of the H&E stain was less intense for the tissue engineered fibrous cap models in comparison to the human cap control (Fig.5A,B). Collagen specific stainings (Fig.5C) further revealed the presence of collagen type III and I of which the latter appeared to have a predominantly increased expression in the C-strain group when compared to the I-strain and predominantly the static group. An increased number of cells are present at the luminal side of the fibrous cap, which is attributable to the presence of an endothelial layer (Fig.5C&D) that is deliberately left-out in our models.

**Figure 5.**
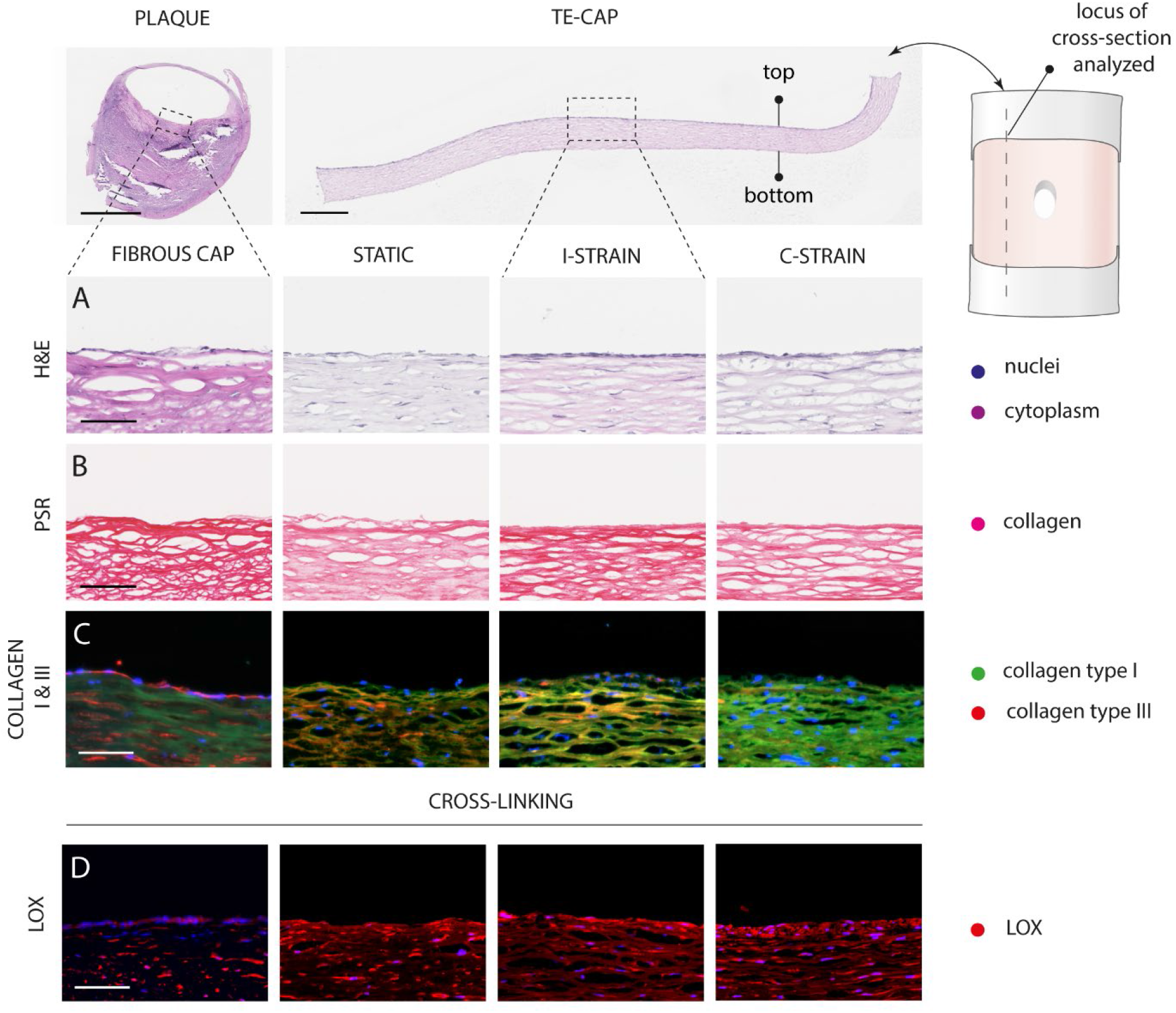
Microscopical evaluation of engineered tissue content at day 21. Cross sections (7 μm) stained for hematoxylin and eosin indicating the cell nuclei (**blue**) and cytoplasm (purple) **(A)**, Picrosirius red (PSR) indicating collagen presence (pink) **(B)**, collagen type I (green) and type III (red) **(C)** lysyl oxidase (LOX, red) **(D)** and cell nuclei (blue) **(C,D)**. Scale bars, 100 μm.

In correspondence to the human cap, LOX was detected in all engineered tissues demonstrating a homogeneous presence of LOX throughout the cross-section (Fig.5D).

### Engineered tissues approach physiological relevant plaque mechanics

Uniaxial tensile tests revealed that all engineered matrices demonstrated physiological strain stiffening responses (Fig.6A) with values that mimic plaque mechanical properties *ex vivo*. More specifically, stiffness values were 0.3 ± 0.2 (static), 0.3 ± 0.1 (I-strain) and 0.4 ± 0.2 (C-strain) MPa at 2% y-strain (ε_yy_),1.1 ± 0.5 (static) 1.1 ± 0.5 (I-strain) and 1.2 ± 0.4 (C-strain) MPa at 5% ε_yy_, and 2.3 ± 0.9 (static), 1.6 ± 0.3 (I-strain) and 2.2 ± 0.9 (C-strain) MPa at 10% ε_yy_ (Fig.6B).

**Figure 6.**
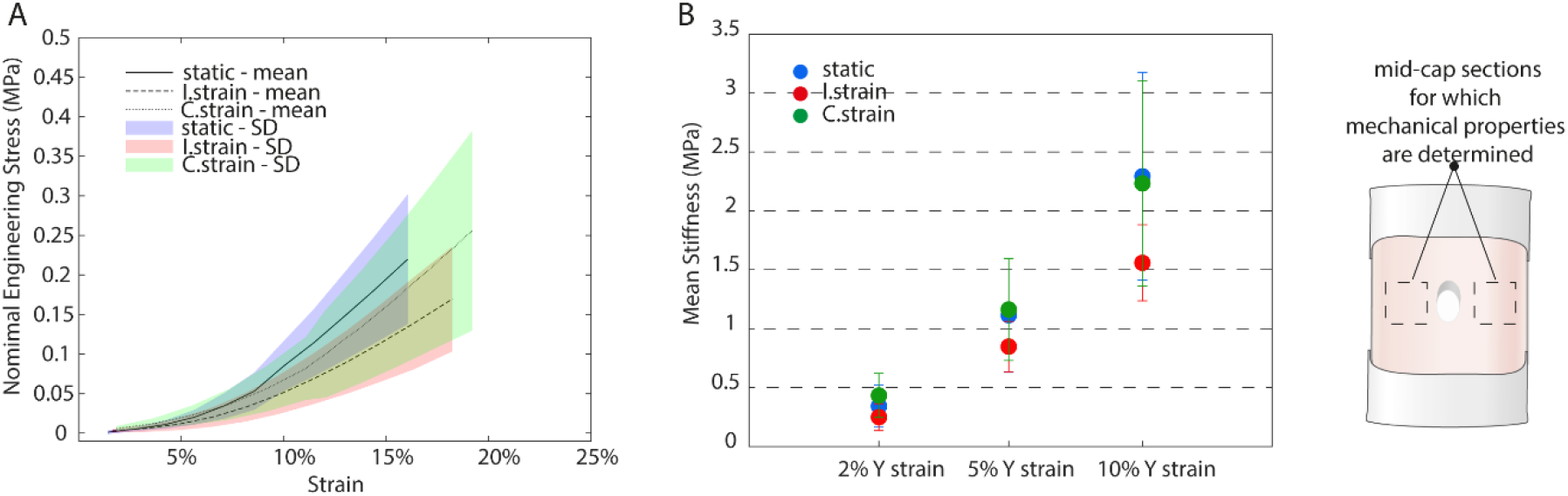
Mechanical characterization of the tissues after 21 days of (dynamic) cell culture. Illustrative nominal engineering stress-strain curves for each experimental group investigated **(A)**. Average calculated stiffness values for the mid cap sections at 2%, 5% and 10% ε_yy_ per experimental group (n ≥ 5 per group) **(B)**.

## Discussion

Within this study we present three tissue engineered fibrous cap models, suitable for mechanical testing, that can each serve as a tunable human in vitro platform to elucidate the reciprocal relationships between atherosclerotic cap composition and mechanics. In contrast to real plaques, the created reductionist models comprise the main attributes that determine cap collagenous matrix mechanical behavior and strength in a controlled manner. All three variants presented show an integrated soft inclusion (i.e. lipid core analog) sealed by a collagen type I and III-positive, organized and crosslinked collagenous ‘cap’ that mimics the microscopic architecture and mechanical behavior of human caps. Importantly, by changing the cell culture conditions applied collagenous matrix content and organization can be tuned. The models allow to properly define global and local tissue properties and can be seen as a first step to elucidate cap failure in vitro.

To create the models dynamic cell culture protocols were selected and compared to static control strategies to create matrices in relative short culture periods with variable collagen content (i.e. amount, type and crosslinking) and organization. Confirming expectations, we observed that cyclic strain, predominantly continuous straining protocol, induced HVSC proliferation ^56–60^. Although cyclic strain is often associated with accelerated matrix deposition, as specifically shown for intermittently strained HVSCs^19^, no difference in total collagen content was detected between the different experimental groups in our study. This might be attributed to the duration of load exposure (i.e. 7 days) as it was previously reported that dynamically loaded HVSCs initially, during the early phases of cell culture, favor a proliferative over a synthetic response ^19,48^.

We obtained more compacted dog-bone shape matrices upon dynamic loading, with no preservation of the initial fibrin gel volume, suggesting enhanced matrix remodeling. Indeed, when evaluating the intrinsic collagenous architecture, clear anisotropic collagen configurations, in the direction of loading, were observed throughout the complete tissue thickness for the dynamically loaded samples as well as in the top layer of the static control. These matrix (re-)orientation responses are known to be a direct consequence of cellular (re-)orientations due to tissue constraints, contact guidance of the endogenous matrix and the magnitude and type of applied loads ^43,45,47,61,62^. Within this study, the Velcro constraints were the main directors of observed collagen anisotropy. Confirming previous studies, it was observed that cells oriented and deposited collagen parallel to the direction of the Velcro constraints ^45^. Moreover, remodeling processes were significantly faster upon dynamic loading but respectively slower in deeper tissue layers in comparison to the superficial top layer due to the contact guidance provided by the endogenous matrix ^45^. As the Velcro constraints were only placed in two positions, in the direction of loading during cell culture, the physiological strain shielding response of HVSCs (i.e. orientating themselves perpendicular to the direction of loading to shield themselves from excessive straining) was prevented. Interestingly, a predominantly isotropic collagen organization was detected in the top and base regions of the dynamically stimulated tissues (i.e. the area between the Velcro constraint and the soft inclusion). These results illustrate that cellular (re-)orientation processes and consequent collagen fiber (re)distributions depend on the cumulative presence of constraints, matrix stiffness and local strain patterns.

Histological comparisons between native fibrous caps and the engineered fibrous cap regions demonstrate that both collagen type I and III, the main load bearing fibrillar collagen types of the fibrous cap ^21–23^, were present in the engineered tissues with qualitative increasing rates of collagen type I upon continuous cyclic loading during cell culture. Engineered fibrous cap collagen type III expression was limited when compared to the human cap control. However, both decreases^63^ as well as increases^64^ in the proportions of type III to I have been observed in atherosclerotic lesions before. Variations in collagen type have been correlated with tissue mechanics^65,66^, though so far no correlations were found between collagen type proportion and fracture stress in fibrous caps^67^Changing cell culture loading protocol offers the opportunity to adapt this ratio, embracing the variance observed in vivo.

Besides collagen content, organization and type we assessed LOX expression because of its catalyzing role in collagen crosslinking. Previous observations revealed a dominant role for collagen crosslinks over collagen content with respect to tissue mechanics in comparable tissue-engineered matrices^17^, bone^68^, scar tissue of collateral ligaments^69^, patellar tendon^70^ as well as cardiovascular tissues^17^. Within this study, we found similar levels of LOX expression between the different tissue engineered fibrous cap models and the human cap control. It is possible that subtle differences in collagenous matrix organization and collagen type are reflected in tissue mechanics. Yet, in line with the similar rate of LOX expression, no statistically significant differences in the global mechanical properties (i.e. elastic moduli) were detected between the dynamically and statically cultured tissues. Although it is difficult to make a direct comparison to human caps, because of their complex composition and the variety of mechanical analyses performed in literature ^6^, all engineered tissues analyzed demonstrated physiological strain stiffening behavior and calculated stiffness values fell within the range reported for human carotid fibrous caps (i.e. 0.5-5 MPa) ^6^. We chose to exclude ultimate tensile strengths values, as many samples, especially the statically cultured, ruptured outside the locus of interest due to local stress accumulations at the Velcro side, underlining the necessity to develop new clamps that improve load transmission. Although a thorough evaluation of local strain patterns is beyond the scope of this study, DIC analyses before rupture demonstrate regions of elevated extension left and right of the soft inclusion. Those regions coincided with the locus of failure for those samples that failed in the region of interest (i.e. the fibrous cap region) for all experimental groups assessed (supplementary Fig.S6).

Finally, we make the note that we made some well-considered choices. First, we chose for a cellular approach because of the inherent advantage that we generate more physiological-like cell made and crosslinked collagenous matrices. Theretofore, cells of the human vena saphena magna were selected as consistent myofibroblastic cell source. They were selected based on their proven role in collagen deposition and remodeling^49^, our previously obtained experience using these cells for cardiovascular tissue engineering purposes^17,19,42,45,62,71^, and the anticipated large number of caps that can be consistently created *in vitro* with cells of a single donor. The human vena saphena magna is often used for coronary bypass surgeries^72^ and adventitial myofibroblast as well as phenotypically altered synthetic VSMCs are deemed responsible for fibrous cap formation ^73,74^. Although comparable cell responses have been observed in VSMCs and between various HVSC donors, potential cell type and donor to donor related variations in matrix deposition, possibly as a result of age, sex and co-morbidity ^75–77^, should be taken into consideration. Creating more personalized models could further enhance risk of rupture stratification.

Second, as human caps develop and adapt over decades and collagen half-life in atherosclerosis might decrease up to 10-fold^78^ due to SMC scarcity and quiescence, it is unrealistic to expect (and as such not our purpose to reproduce) an exact mimic. The purpose of our *in vitro* model is to create a physiological-like tunable platform that will be iteratively used, together with more simplified *in vitro* and more complex *ex vivo* models (e.g. human plaque lesions), to test, validate and revise hypotheses on matrix mechanics and failure. Importantly, the model is suitable for high frequency and resolution real-time imaging to study collagen evolution and failure instantaneously. By elongating load duration or changing load type, it is expected that the engineered collagenous tissue properties as demonstrated in this study can be further manipulated^17,19,42,45,71^. To stretch the boundaries of our current approach, additives like TGF β^79^, ribose^80,81^, copper^82^ or beta-aminopropionitrile^83^ might be added, to further stimulate collagen deposition, induce AGE related crosslinking and enhance or inhibit LOX catalyzed crosslinking respectively.

Lastly, plaque and cap rupture cannot be attributed to solely suboptimal collagenous properties. It is a cumulative consequence of these properties, local inflammatory cell driven collagen degradation, and steep gradients in stiffnesses and loading resulting in local stress accumulations ^84–89^. The current model can be used as a first step covering cap complexity. Stiff calcifications, intraplaque hemorrhages, inflammatory cells or substitutes can be added to further embrace physiological variance and plaque failure modes. Overall, the model will aid in understanding material properties, local strain fingerprints, stress accumulations and tissue failure modes in collagenous tissues. Information that can be implemented in *in silico* models to improve biomechanical stress analyses and the assessment of plaque vulnerability specifically or tissue damage in complex heterogeneous tissues in general ^6^.

## Conclusion

To conclude, we present three tissue-engineered fibrous cap models that aid in the controlled and systematic investigation of the reciprocal relationships between cap composition and mechanical properties. By applying different (dynamic) cell culture protocols we were able to create reproducible naturally crosslinked tissues, with a tunable collagen content and organization, suitable for mechanical testing. All models contained a successfully integrated soft inclusion (i.e. lipid core analog) that was covered by a collagen type I and III-positive organized ‘cap’, to simulate plaque tissue heterogeneity. These caps exhibited physiological strain stiffening responses and calculated stiffness values that fell within the ranges reported for human caps in literature. Each model can be seen as a tunable physiological-like 3D platform, that can be iteratively used, together with more simplified *in vitro* models and more complex *ex vivo* material, to ultimately study and elucidate cap rupture. However, the model can also be of value to understand tissue mechanics and damage in complex heterogeneous tissues in general. Moreover, as this model is suitable for high-resolution, high-frequency real-time imaging, it can be used to investigate collagen remodeling and failure at the microlevel. Future efforts should focus on integrating stiff inclusions, potentially calcifications, or collagen degrading factors, like macrophages, to further aid in the identification of cap rupture risk.

## Acknowledgements

We gratefully acknowledge the Gravitation Program “Materials Driven Regeneration”, funded by the Netherlands Organization for Scientific Research (024.003.013) and Kim van Gaalen, Sylvia Dekker and Mark van Turnhout for their contribution with the analyses. Kim van der Heiden is funded by an NWO-Vidi grant 18360.

## Conflict of Interest

The authors declare no conflict of interest.

## Notes

### Competing Interest Statement

The authors have declared no competing interest.

