## supplementary information for "Tissue-engineered collagenous fibrous cap models to systematically elucidate atherosclerotic plaque rupture"

1 Department of Biomedical Engineering – Thorax Center Erasmus Medical Centre, Rotterdam, The Netherlands, 2 Department of Biomedical Engineering, Eindhoven University of Technology, Eindhoven, the Netherlands 3 Institute for Complex Molecular Systems (ICMS), Eindhoven University of Technology, Eindhoven, The Netherlands, 4 Biomechanical Engineering, Technical University Delft, Delft, The Netherlands.

Department of Biomedical Engineering, Biomechanics Laboratory Ee2341, P.O. Box 2040, 3000 CA Rotterdam, The Netherlands. Tel.: +31 10 704 4045; fax: +31 10 704 4720*

| Supp. Table S1. Average Y-strains applied during dynamic cell culture– day 21 | | |
| --- | --- | --- |
|  | **Average Y-strain ± SD (%)** | **n** |
| I-strain (Exp 1) | 4.3 ± 0.0 % | 2 |
| C-strain (Exp 1) | 7.8 ± 0.8 % | 3 |
| I-strain (Exp 2) | 5.3 ± 0.3 % | 4 |
| C-strain (Exp 2) | 5.1 ± 0.7 % | 4 |
| I-strain (Exp 1+2) | 4.9 ± 0.5 | 6 |
| C-strain (Exp 1+2) | 6.3 ± 1.6 | 7 |

The average maximum Y-strains (direction of loading) during culture in the region of interest (see supplementary S1) were determined by correlation (DIC) analysis at day 21. On average, samples of the I-strain group were exposed to 4.9 ± 0.5 % strain where the C-strain samples were subjected to a slightly higher and more variable 6.3 ± 1.6 % strain. This variation is predominantly caused by the difference in loading during the first of the two experiments performed; i.e. 4.3 ± 0.0 % I-strain vs. 7.8 ± 0.8 % C-strain (Exp1) and 5.3 ± 0.3 % I-strain vs. 5.1 ± 0.7 % C-strain (Exp 2), respectively. These variations in loading didn’t affect obtained collagen content and mechanical properties.

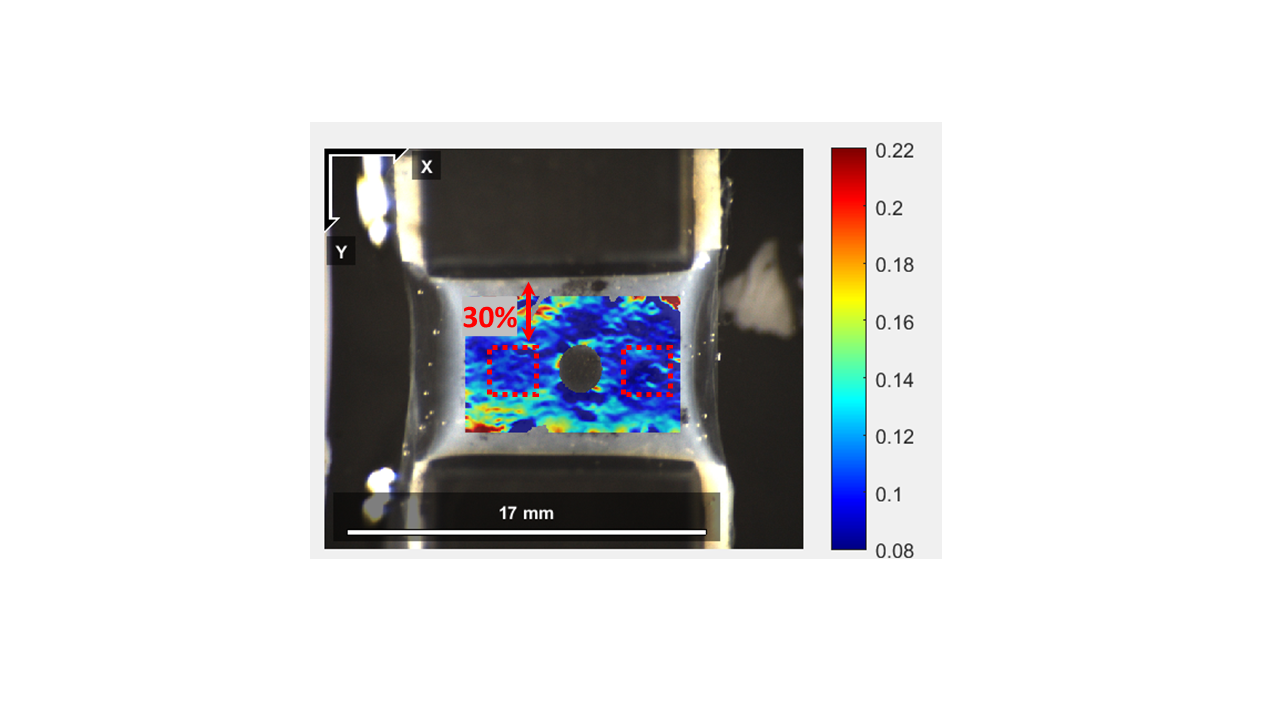

**Supp. Fig. S1.** Regions of interest (ROI, indicated by the red dotted lines) for which DIC analyses were performed. 30 percent from the edges were excluded to prevent that compaction and clamping artifacts were included in the analyses. The regions on each side of the soft inclusion were treated separately as local variation may occur. The average strain was taken of both regions of interest per construct.

**Method for determining fiber organization**

The mean predominant fiber angle for each region (mid cap, shoulder, top) was determined for each group (static, I-strain, C-strain) by capturing CNA stained fluorescent microscopy images per region and inputting them into a MATLAB based fiber orientation analysis tool previously developed [1].

Images were captured of both the top and bottom of the tissue to get representative images of the condensed ± 5 – 50 µm thick top surface layer (1 - 10% of the tissue thickness) and the remainder ≥ 90 percent of the tissue thickness that displayed consistent organizational patterns throughout its depth.

Images were captured from the top and bottom at approx. 15 μm steps in depth until approx. 60 μm depth. Due to signal attenuation with increased depth, one image per region was chosen for fiber orientation analysis in both the top (first 5 - 50 μm) and bottom surface layers (S.Fig 1).

0 μm

**TOP**

30 μm

Depth

Tissue thickness

60 μm

60 μm

Depth

30 μm

**BOTTOM**

0 μm

**Supp. Fig. S2.** Cross sectional view of tissue. The images selected for fiber orientation analysis were within the pink region (depth between 15-30 μm); not to scale.

A Gaussian distribution was fit to the histogram orientation data of each image using FibLab, an in-house MATLAB based tool developed at TU/e for fitting angle distributions [2]. FibLab considers the data located above the ‘isotropic baseline’, *b*, as exhibiting anisotropic behavior and uses only this data for the Gaussian fitting. The peak angle(s), μ_p_, independent of the baseline, as well as the standard deviation value(s) per peak, σ_p_, are reported (Fig 2). The σ_p_ is representative of the degree of anisotropic behavior along a predominant direction (μ_p_). In cases where ‘wrapping’ occurs at the edges (near 0° and 180°), FibLab treats the data as a continuous curve and reports one peak angle and standard deviation value (Fig 3.a). Figure 3 represents the most common distributions encountered in this experiment and how FibLab determined μ_p_ and σ_p_ for each case.

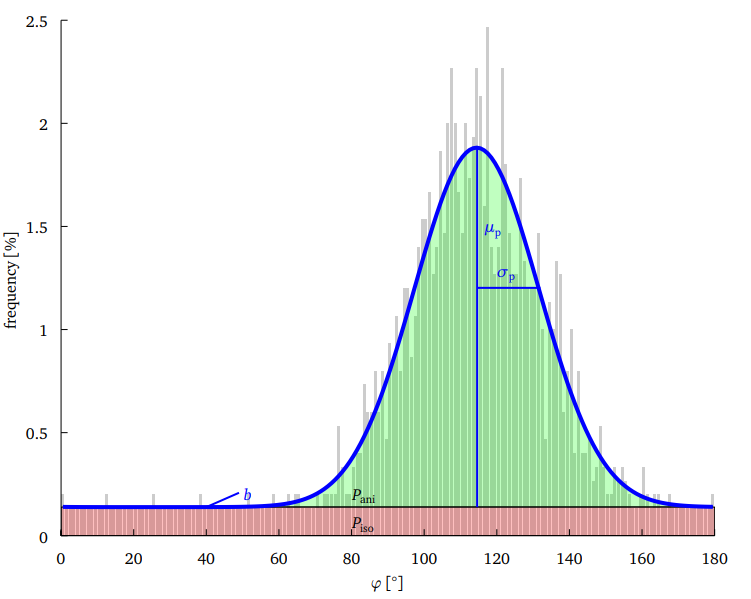

**Supp. Fig.S3.** Arbitrary distribution example demonstrating the values reported in FibLab: μ_p_, the center of the peak (‘mean’ of the fitted normal distribution); σ_p_, the standard deviation of the peak; *b*, baseline [2]

(a)
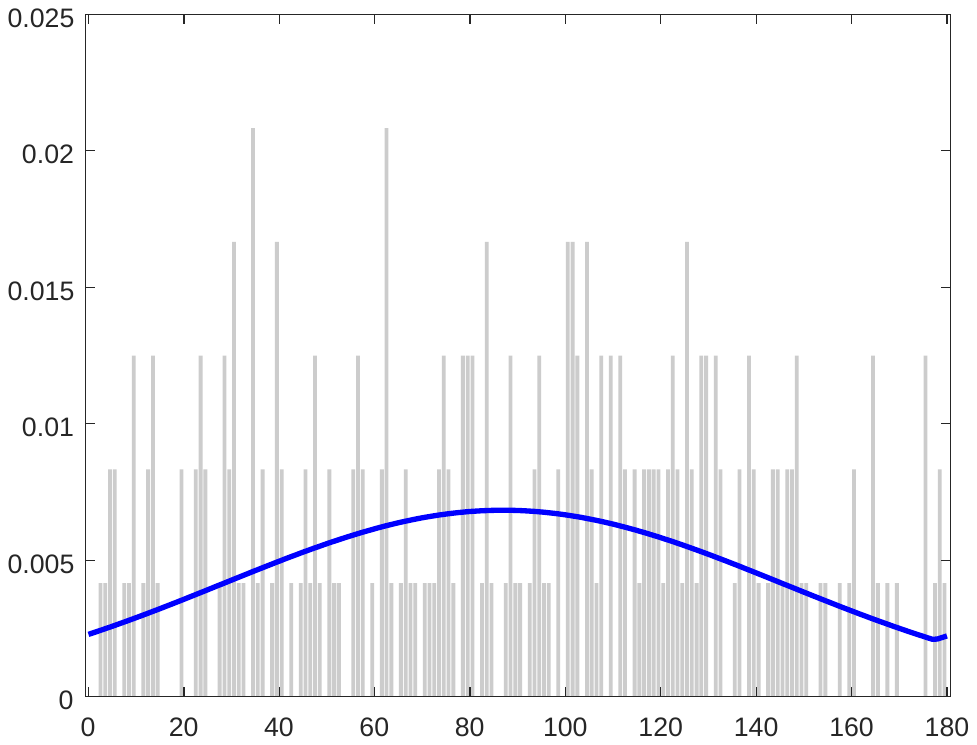

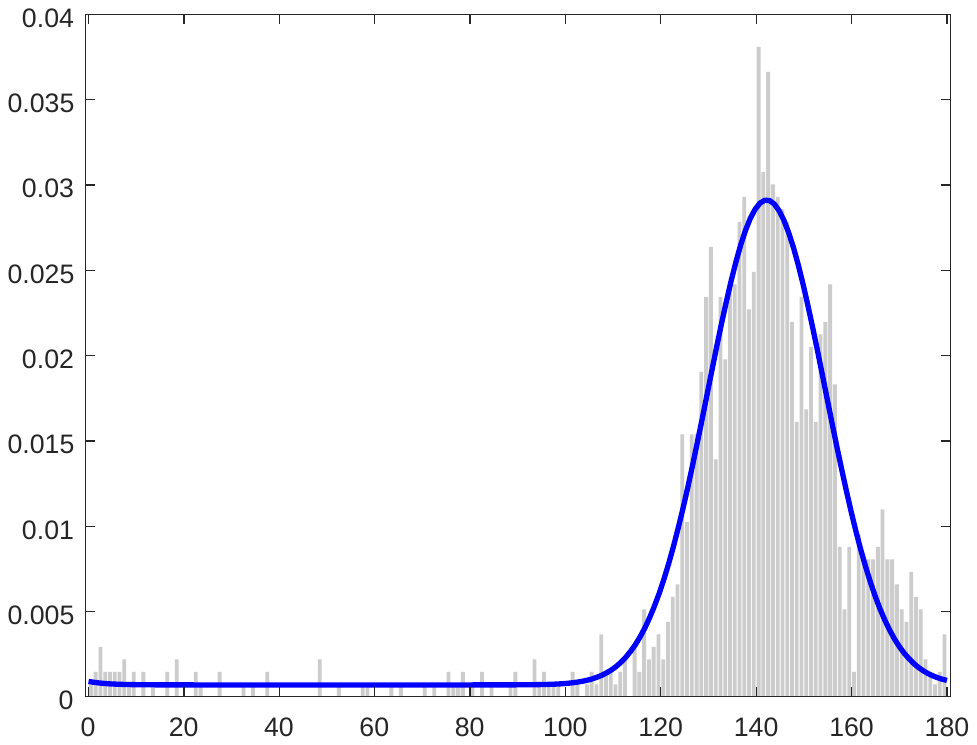

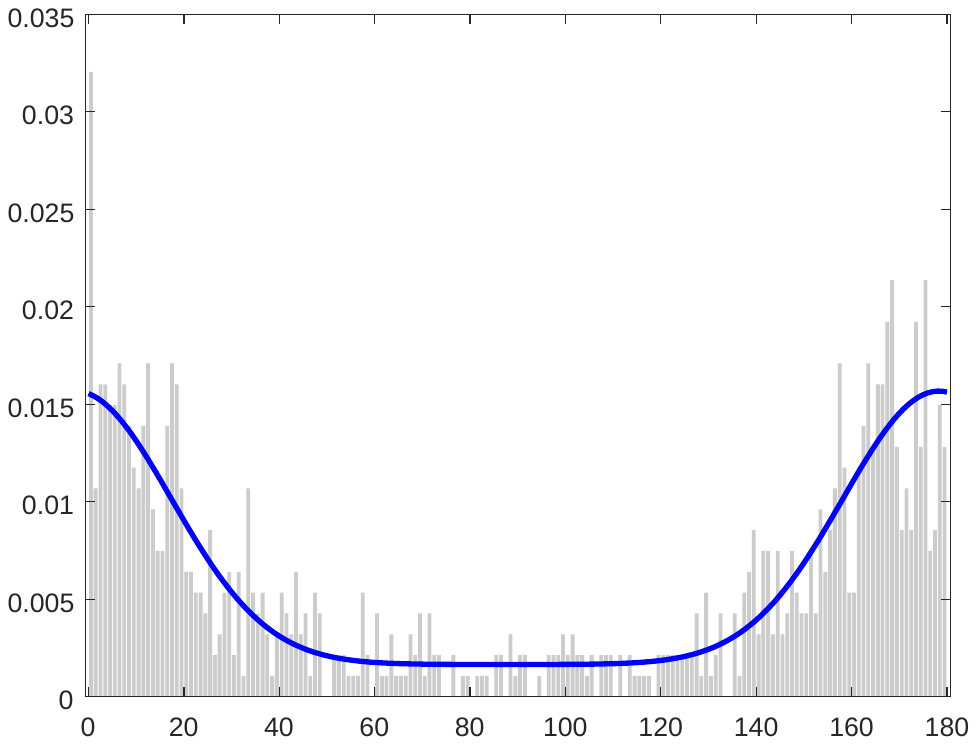
 (b)	(c)

**Supp. Fig.S4.** Typical examples of the fiber angle data distributions in this experiment, Gaussian fit (blue): (a) two wrapping peaks, FibLab considers this data as one continuous peak μ_p_ = 178°, σ_p_ = 20° (b) one peak, μ_p_ = 142°, σ_p =_ 12° (c) one wide peak (demonstrates more isotropic behavior, μ_p_ = 87°, σ_p_ = 58°)

In order to simplify whether the fibers were oriented parallel or perpendicular to the loading direction, the resulting predominant fiber angles larger than 90° (∠_>90_) were converted from FibLab’s default 0-180° scale to a 0-90° scale (∠), where 0° represents the loading direction (Eq. 1). The mean predominant fiber angles ± mean σ_p_ for each region per straining group were calculated and plotted (Fig # in paper)

∠ = 180 - ∠_>90_	**Supp. Eq. 1**

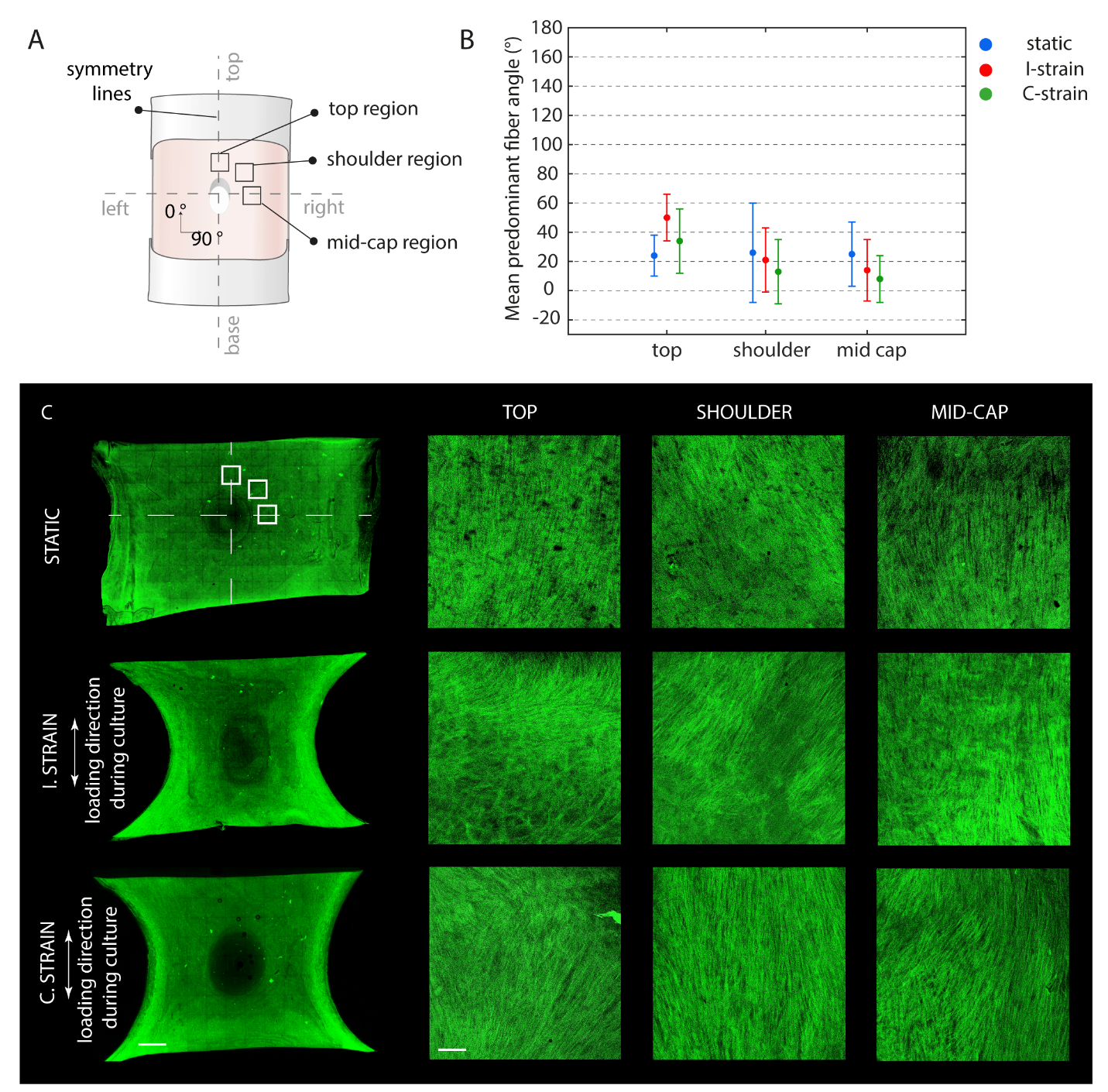

**Supp. Fig.S5** Collagen architecture of the condensed ± 5 – 50 µm thick upper superficial layer (1 - 10% of the tissue thickness) at day 21. Graphical representation of the loci visualized **(A).** Quantification of the collagen fiber distribution (i.e. predominant fiber angles and presence of (an)isotropy). Large SD indicates greater isotropic behavior surrounding mean predominant fiber angle. Small SD indicates greater anisotropic behavior surrounding mean predominant fiber angle **(B)**. Representative confocal tile scans and zoom-ins of CNA35 (a collagen binding protein ^53^, green) stained samples of the same top, shoulder and mid-cap regions as evaluated in B **(C)**. Scale bars, 3 mm (tile scans) and 100 µm (zooms).

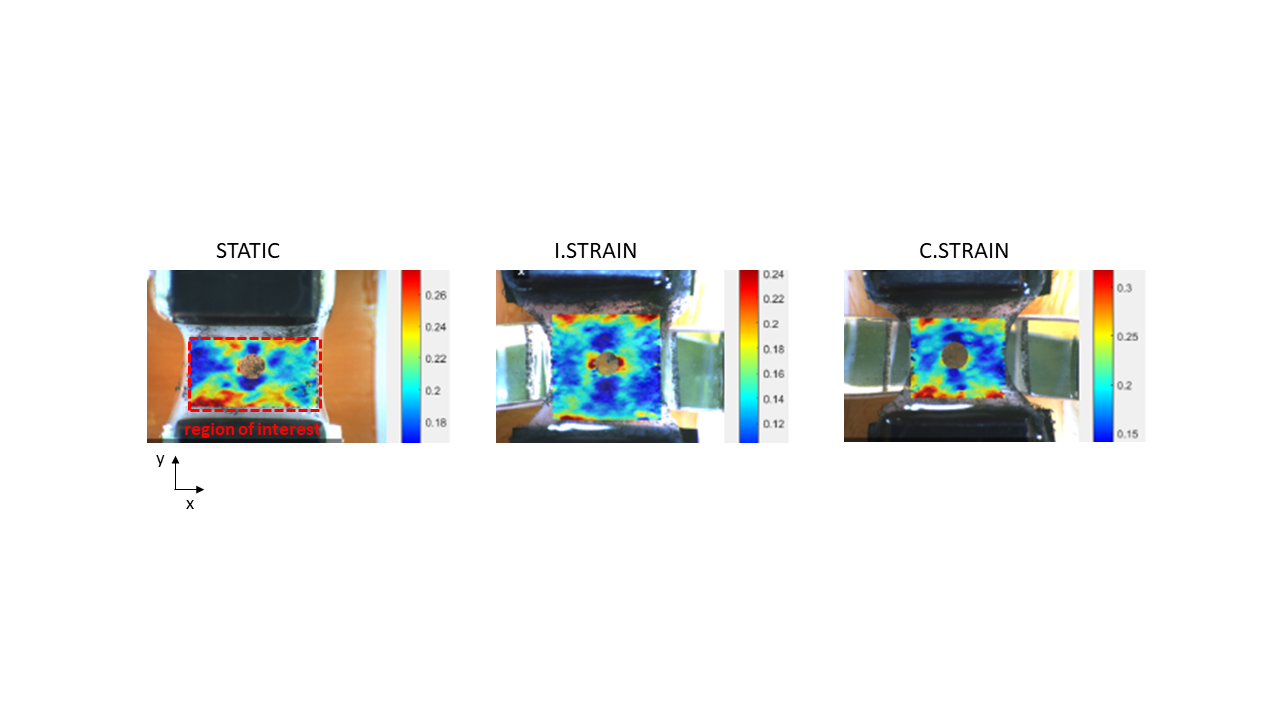

**Supp. Fig.S6** DIC analyses before rupture demonstrate the highest strains in the direction of the loading around the clamp edges and left and right of the soft inclusion for all groups analyzed. To prevent the tissue from failing outside our region of interest (red dotted line), at the clamp edge, new clamps should be developed that improve load transmission.
